# TBACdb: a comprehensive database of tobacco bioactive compounds

**DOI:** 10.64898/2026.11.20.767781

**Authors:** Wentao Cui, Runtong Hou, Zhican Fu, Xuejie Zhang, Rui Nie, Jingcheng Zhang, Xinkui Zhao, Shaofeng Liu, Peng Lu, Yong Chen, Tietuo Tao, Junlan Cai

## Abstract

Tobacco (*Nicotiana tabacum* L.) contains diverse chemical constituents, but information on their biological activities, plant origins, supporting literature, and experimentally accessible standards remains fragmented. Here, we developed TBACdb (http://115.120.251.12:30003/), an integrated database of tobacco bioactive compounds. Starting from 5,139 tobacco-associated chemicals with CAS numbers, multi-source curation yielded 4,634 standardized compound records, including 2,377 compounds associated with reported biological activities and 166 compounds directly supported by tobacco-focused studies. TBACdb integrates 64,522 compound–bioactivity records, 12,465 compound–species associations, more than 3.7 million literature records, and 783 physical reference standards. The database provides chemical structures and physicochemical properties, biological activities, species origins, literature evidence, statistical analysis, external database links, and digital–physical compound mapping. TBACdb provides a centralized resource for tobacco compound exploration, candidate prioritization, and subsequent analytical and experimental investigation.

## 1 Background

Tobacco (*Nicotiana tabacum* L.) is an economically important plant characterized by a diverse array of primary and specialized metabolites. Plant specialized metabolites play important roles in ecological interactions and represent a valuable source of structurally diverse natural products [1**?** –3]. Tobacco contains numerous chemical constituents, including alkaloids, terpenoids, phenolic compounds, flavonoids, and organic acids, many of which have attracted attention for their biological properties and potential applications [4–6]. Representative compounds such as nicotine, solanesol, chlorogenic acid, and rutin have been investigated in various biological and pharmacological contexts [7–10]. Despite these advances, available studies have primarily focused on selected well-characterized compounds, while comprehensive information on the chemical diversity, biological activities, and experimental resources of tobacco constituents remains fragmented.

The development of chemical and natural-product databases has substantially facilitated the systematic organization and exploration of compound-related information. PubChem provides extensive chemical structures, physicochemical properties, biological activities, and literature information [11, 12]. Natural-product databases such as NPASS and CMAUP integrate biological activities, molecular targets, and plantsource information [13, 14], whereas KNApSAcK focuses on relationships between plant species and their metabolites [15]. Other resources, including COCONUT and LOTUS, provide large-scale collections of natural products and their reported biological origins [16, 17]. In addition, ChEBI and NPClassifier support standardized chemical annotation and structural classification [18]. Although these resources have significantly advanced natural-product research, they are primarily general-purpose databases rather than integrated resources specifically designed for tobacco-associated bioactive compounds.

Several challenges remain in the systematic investigation of tobacco bioactive compounds. First, chemical properties, biological activities, species origins, and supporting literature are distributed across heterogeneous resources, making cross-database integration and compound-level analysis difficult [19]. Second, biological activities reported for the same chemical entity in different plant species do not necessarily constitute direct experimental evidence from tobacco-derived materials. Distinguishing tobaccospecific evidence from broader plant-derived annotations is therefore important for reliable interpretation and candidate prioritization. Third, existing digital resources generally provide limited integration with physical reference materials, which are essential for compound identification, quantitative analysis, and experimental validation. A tobacco-centered resource combining standardized chemical information, traceable biological evidence, and experimentally accessible reference standards would help address these limitations.

To address these needs, we developed the Tobacco Bioactive Compounds Database (TBACdb; http://115.120.251.12:30003/), an integrated resource connecting tobaccoassociated chemical compounds, biological activities, species origins, literature evidence, and physical reference standards. TBACdb contains 4,634 standardized tobacco compounds, including 2,377 compounds associated with reported biological activities, together with 64,522 compound–bioactivity records, 12,465 compound–species associations, and more than 3.7 million literature records. In addition, 783 compounds are linked to corresponding physical reference standards. By organizing heterogeneous information within a unified compound-centered framework, TBACdb supports systematic compound exploration, evidence tracing, candidate screening, and subsequent experimental investigation.

## 2 Construction and content

### 2.1 Data sources and collection

TBACdb integrates tobacco chemical information, biological activities, species origins, literature evidence, and physical reference materials from multiple heterogeneous sources. The initial compound set was compiled from authoritative references on tobacco and tobacco-smoke chemistry [20], yielding 5,139 tobacco-associated compounds with available CAS registry numbers. Chemical structures and physicochemical properties were collected primarily from PubChem [11] and supplemented with DrugBank [21], ChemicalBook, JheChem, Reaxys, ChemSpider, COCONUT [16], and related chemical resources. These sources provided complementary information on compound names and synonyms, molecular formulas, molecular weights, SMILES, InChI, InChIKey, structural representations, and physicochemical properties. Biological activity and species-origin information was obtained from natural-product and plant-oriented resources, including NPASS [13], CMAUP [14], PlantPlus, pharmacopoeias, reference books, and relevant scientific literature. Literature information was collected mainly from PubMed, ScienceDirect, CNKI, Wanfang, and related literature databases. To distinguish tobacco-specific evidence from broader functional information, publications directly investigating tobacco-derived compounds were manually screened, resulting in 166 compounds with directly reported biological activities, whereas cross-referencing the tobacco compound set with broader plant-derived resources expanded functional annotations to 2,377 compounds. Data acquisition combined database interfaces, automated retrieval, batch processing, document parsing, and manual curation. For document-based resources, text extraction and optical character recognition were applied when necessary, followed by manual verification of ambiguous compound names, identifiers, and activity descriptions. Source information and original identifiers were retained during data integration to support provenance tracing, quality control, and subsequent updating. The major data sources and information types incorporated into TBACdb are summarized in Table 1.

**Table 1:**
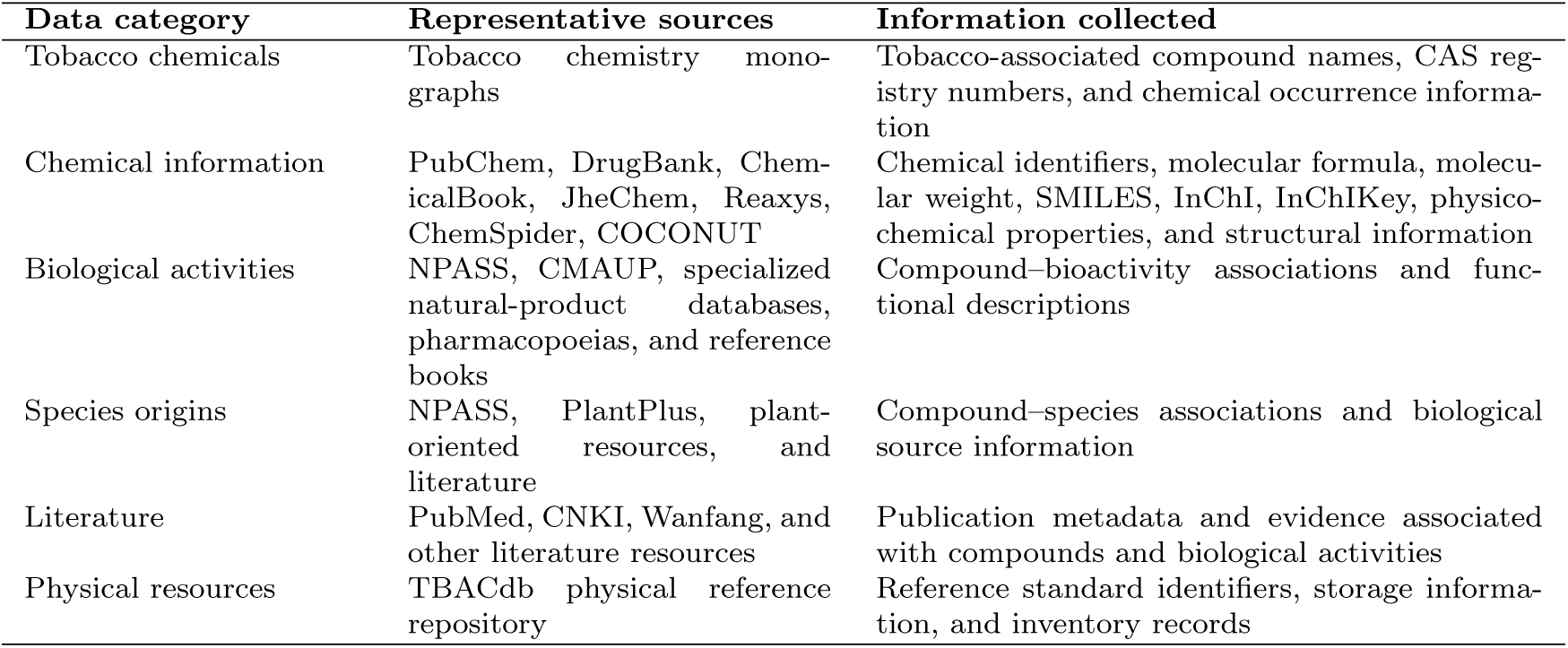
Major data sources integrated into TBACdb.

| **Data category** | **Representative sources** | **Information collected** |
|---|---|---|
| Tobacco chemicals | Tobacco chemistry mono-graphs | Tobacco-associated compound names, CAS reg-istry numbers, and chemical occurrence informa-tion |
| Chemical information | PubChem, DrugBank, Chem-icalBook, JheChem, Reaxys, ChemSpider, COCONUT | Chemical identifiers, molecular formula, molecu-lar weight, SMILES, InChI, InChIKey, physico-chemical properties, and structural information |
| Biological activities | NPASS, CMAUP, specialized natural-product databases, pharmacopoeias, and reference books | Compound–bioactivity associations and func-tional descriptions |
| Species origins | NPASS, PlantPlus, plant-oriented resources, and literature | Compound–species associations and biological source information |
| Literature | PubMed, CNKI, Wanfang, and other literature resources | Publication metadata and evidence associated with compounds and biological activities |
| Physical resources | TBACdb physical reference repository | Reference standard identifiers, storage informa-tion, and inventory records |

### 2.2 Collection and standardization of tobacco compounds

The initial 5,139 tobacco-associated compounds were subjected to a unified standardization procedure before downstream integration. CAS registry numbers were used as the primary identifiers for cross-database matching, while SMILES, InChI, and InChIKey provided additional structural information for compound identity verification [22]. PubChem served as the principal source of standardized chemical information [11], including preferred names, molecular properties, and structural representations, whereas DrugBank, ChemicalBook, JheChem, Reaxys, COCONUT, and other resources were used to supplement missing attributes and resolve incomplete or inconsistent records. Records referring to the same chemical entity were merged on the basis of concordant identifiers and structural information, while alternative names, synonyms, and source-specific identifiers were retained to facilitate traceability and cross-resource retrieval. When discrepancies were observed among databases, standardized records were determined by cross-comparing available identifiers and structural descriptors, with priority given to more authoritative and complete sources. After identifier matching, duplicate removal, consistency checking, and manual review of ambiguous records, 4,634 standardized tobacco compounds were retained in the current release of TBACdb. Chemical classification was further integrated from COCONUT [16] and supplemented using NPClassifier, enabling compounds to be organized into hierarchical natural-product categories. These standardized compound entities constitute the central backbone of TBACdb and provide unified links to biological activities, species origins, literature evidence, and physical reference materials (Fig. 1).

**Fig. 1:**
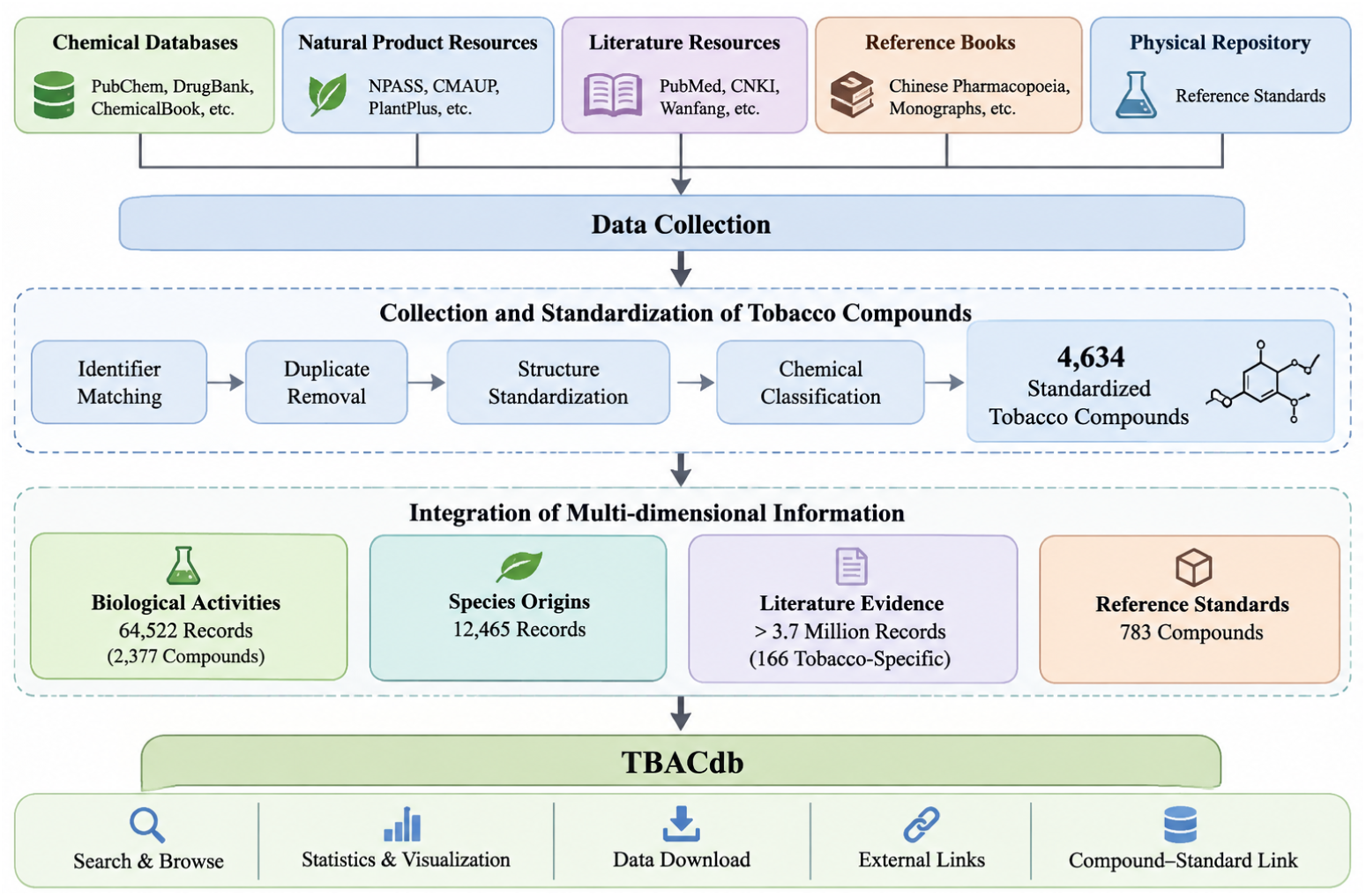
Overview of the data collection, standardization, integration, and functional organization of TBACdb. Tobacco-associated compounds were collected from chemical databases, natural-product resources, literature databases, reference books, and the physical reference repository. Compound records were standardized through identifier matching, duplicate removal, structure standardization, and chemical classification. The resulting standardized compounds were further linked to biological activities, species origins, literature evidence, and physical reference standards, forming the integrated TBACdb resource.

### 2.3 Integration and curation of biological activity information

Biological activity information was integrated through two complementary routes to distinguish tobacco-specific evidence from broader functional evidence reported for the same chemical entities in other plant resources. First, tobacco-focused publications were manually screened, and reported compounds were matched to the standardized tobacco compound set primarily using CAS registry numbers, with compound names and structural identifiers used when necessary. Biological functions and activity descriptions were extracted and linked to the corresponding standardized records, resulting in 166 compounds with biological activities directly supported by studies of tobacco-derived materials. Second, the standardized tobacco compounds were cross-referenced with natural-product databases, pharmacopoeias, reference books, and related plant-derived resources, including NPASS and CMAUP [13, 14]. When an identical chemical entity was reported in another plant resource, its associated biological activity information was incorporated as complementary functional evidence, expanding the coverage to 2,377 compounds with reported biological activities. Activity descriptions collected from heterogeneous sources were subsequently normalized to reduce redundant or inconsistent annotations, and duplicate compound–activity associations were consolidated where appropriate. After integration and curation, TBACdb contained 64,522 compound–bioactivity records. Importantly, activity information obtained through cross-plant matching was retained separately from evidence directly derived from tobacco-focused studies, because a biological effect reported for the same compound in another plant does not necessarily indicate experimental validation in tobacco-derived material. Source information and supporting literature were therefore preserved whenever available, allowing users to trace the provenance and interpret the evidence context of individual activity annotations.

### 2.4 Integration of species origins and literature evidence

Species-origin information was integrated to provide biological provenance and distribution context for tobacco-associated compounds. Records were collected from natural-product and plant-oriented resources, including NPASS [13], PlantPlus, and complementary scientific literature, and were mapped to the standardized tobacco compound set using CAS registry numbers, normalized compound names, synonyms, and other available chemical identifiers. After identity matching, normalization, and deduplication, 12,465 compound–species associations were retained in TBACdb. Because an individual compound may occur in multiple plant species and each species may contain multiple compounds, these relationships were represented as many-tomany associations, with original source information preserved for provenance tracing. Literature information was collected mainly from PubMed, CNKI, Wanfang, and other scientific literature resources and linked to standardized compounds using compound names, synonyms, chemical identifiers, tobacco-related terms, and relevant biological descriptors. More than 3.7 million literature associations are currently indexed in TBACdb. These records provide broad bibliographic context rather than uniformly manually curated experimental evidence; therefore, publications directly investigating tobacco-derived compounds were distinguished from broader literature associated with the same chemical entities. By integrating species-origin and literature information, TBACdb enables users to trace both the reported biological distribution of individual compounds and the supporting publications associated with their chemical and functional annotations.

### 2.5 Construction of the physical reference repository

A physical reference repository was established to connect the digital information in TBACdb with experimentally accessible materials. Based on the curated tobaccoassociated bioactive compound collection and subsequent analytical needs, a total of 783 reference standards were collected and incorporated into the repository. Each standard was assigned a unified internal identifier and linked to its corresponding standardized digital compound record, enabling consistent mapping between database information and physical materials. The inventory management system records key information, including compound identity, storage location, inventory status, material amount, and circulation records, and supports the major processes of registration, storage, retrieval, lending, return, and inventory monitoring. Reference standards were stored according to their chemical properties and specific storage requirements to support standardized management and long-term preservation. Barcode-based identification, inventory monitoring, and dynamic record updating were further implemented to maintain consistency between the physical repository and the online database. Through this digital–physical linkage, users can determine whether a reference standard is available while examining a compound record, thereby facilitating subsequent compound identification, quantitative analysis, method development, and experimental validation.

### 2.6 Data curation and quality control

A multi-step quality-control procedure was applied to ensure the consistency, reliability, and traceability of TBACdb. CAS registry numbers were used as the primary identifiers for cross-database integration, while compound names, molecular formulas, molecular weights, SMILES, InChI, and InChIKey were used for additional identity and structure verification. Duplicate records were merged, abnormal or conflicting values were examined, and missing attributes were supplemented from reliable alternative resources when available. For records represented in multiple databases, key identifiers and structural information were cross-compared to identify inconsistencies, and more authoritative or complete sources were preferentially used for standardized fields while alternative annotations were retained when useful for traceability. Biological activity, species-origin, and literature records were normalized and deduplicated while preserving their original provenance. In particular, tobacco-specific biological activity evidence was kept distinguishable from annotations obtained by matching the same chemical entities to other plant-derived resources, thereby reducing the risk of interpreting cross-plant evidence as direct validation in tobacco. Version and source information were retained to facilitate subsequent correction and updating. Physical reference materials were linked to standardized digital records using unified identifiers and barcode-based management, and inventory changes were synchronized with corresponding database records to maintain consistency between the digital and physical repositories.The retention of standardized identifiers and provenance information also supports broader principles of scientific data interoperability and reuse [23].

### 2.7 Database architecture and implementation

TBACdb was implemented using a modular four-layer architecture comprising data, service, application, and presentation layers. The data layer stores the core resources of the database, including compound information, biological activities, species origins, literature records, and physical reference materials. Structured data were organized in a MySQL relational database using modular tables and one-to-many or many-tomany relationships, while large files such as molecular structure images and literature documents were managed separately using object storage. Frequently queried fields, including compound names, CAS registry numbers, and other chemical identifiers, were indexed to improve retrieval efficiency. The service layer provides standardized data-access and business-logic functions for compound retrieval, biological activity querying, species-origin exploration, literature association, statistical analysis, user management, and physical-resource management. The application layer integrates these services into user-oriented functions such as keyword search, multi-condition filtering, data browsing, statistical analysis, and reference-standard management. The presentation layer provides a web-based interface for interactive access to TBACdb and supports direct links to external resources including PubChem, PubMed, and DrugBank. The modular architecture separates data management from user-facing functions, facilitating system maintenance, incremental data updating, and future extension of the database.

## 3 Database content and statistics

### 3.1 Overview of TBACdb

The current release of TBACdb provides an integrated resource for the systematic exploration of tobacco-associated chemical compounds and their biological information. Following multi-source collection, standardization, and curation, 4,634 tobacco-associated compounds with standardized chemical information were retained in the database, of which 2,377 are associated with at least one reported biological activity. In total, TBACdb integrates 64,522 compound–bioactivity records and 12,465 compound–species associations, providing functional annotations and biological-source information for a substantial proportion of the standardized compound collection. More than 3.7 million literature records are further associated with these compounds and their related biological information, supporting compound-centered evidence tracing and bibliographic exploration.

In addition to digital information, TBACdb incorporates 783 physical reference standards that are linked to their corresponding standardized compound records. This digital–physical integration enables users to examine chemical properties, biological activities, species origins, and supporting literature while simultaneously determining whether an experimentally accessible reference material is available. The major data components of TBACdb are summarized in Table 2. By organizing heterogeneous resources around standardized compound entities, TBACdb establishes an interconnected framework linking chemical information, biological functions, plant origins, literature evidence, and physical materials, thereby providing the basis for subsequent analyses of chemical diversity, biological activity patterns, species distributions, and experimental accessibility.

**Table 2:** Summary of the major data resources in TBACdb.

| **Data category** | **Number of records** |
|---|---|
| Standardized tobacco chemical com-pounds | 4,634 |
| Compounds associated with biological activities | 2,377 |
| Compound–bioactivity records | 64,522 |
| Compound–species records | 12,465 |
| Associated literature records | >3.7 million |
| Physical reference standards | 783 |

### 3.2 Chemical diversity of tobacco compounds

To characterize the chemical diversity represented in TBACdb, the 4,634 standardized tobacco-associated compounds were organized according to structural and chemical classification annotations. Classification information was primarily integrated from COCONUT [16] and supplemented using NPClassifier, enabling compounds to be described at multiple hierarchical levels, including chemical superclass, chemical class, and natural-product pathway-related categories. The resulting annotations revealed broad structural diversity across the tobacco chemical space, covering alkaloids, terpenoid-related compounds, phenolic compounds, flavonoids, organic acids and their derivatives, lipids and lipid-like molecules, and numerous other natural-product classes. The distribution of major chemical categories is summarized in Fig. 2. In addition to relatively abundant classes, a substantial fraction of compounds was distributed across less frequently represented structural groups, indicating that TBACdb extends beyond a limited set of well-studied tobacco constituents. Chemical classification information was further linked to biological activity annotations, allowing comparison of the proportion and diversity of bioactive compounds across different structural categories. This organization provides a basis for examining potential relationships between chemical structure and reported biological function and can also highlight compound classes that remain comparatively under-characterized. By integrating hierarchical chemical classification with functional annotations, TBACdb enables users to explore tobaccoassociated compounds from both structural and biological perspectives and provides a foundation for subsequent analysis of the biological activity landscape.

**Fig. 2:**
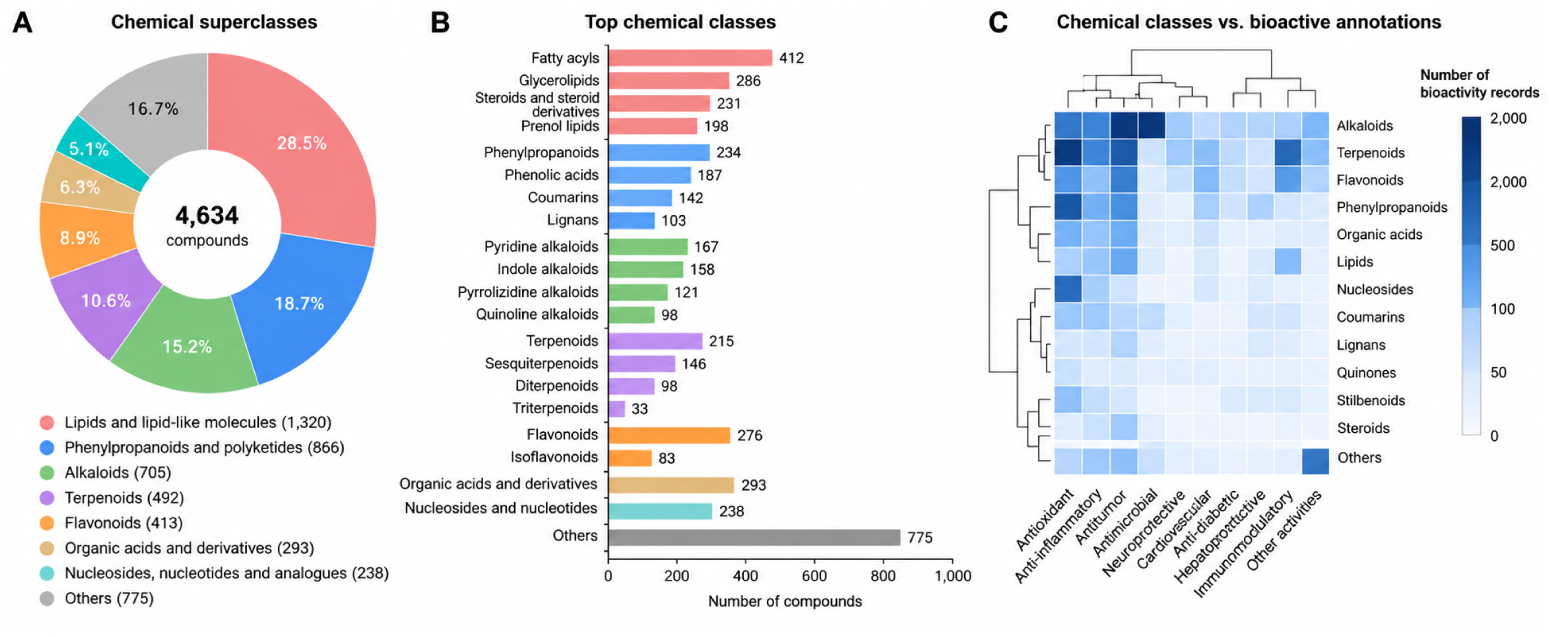
Chemical diversity of compounds included in TBACdb. (A) Distribution of tobacco compounds among major chemical superclasses. (B) Distribution of compounds among major chemical classes. (C) Relationship between chemical classes and bioactive compounds. Only the most abundant categories are displayed, while the remaining categories are grouped as “Others” for visualization.

### 3.3 Landscape of biological activities

To characterize the functional landscape represented in TBACdb, we analyzed the biological activity annotations associated with the standardized tobacco compounds. Among the 4,634 compounds, 2,377 were linked to at least one reported biological activity, corresponding to 64,522 compound–bioactivity records. These annotations were derived from natural-product resources such as NPASS and CMAUP [13, 14], together with reference materials and scientific literature, and cover a broad range of functional categories, including antioxidant, anti-inflammatory, antimicrobial, antitumor, cardiovascular, neuroprotective, hepatoprotective, metabolic, and other reported biological effects. Because individual compounds may be associated with multiple activities and each activity category may involve compounds from different chemical classes, the resulting compound–activity relationships form a many-to-many network. The distribution of major activity categories and their relationships with chemical classes are summarized in Fig. 3. Differences in the numbers of associated compounds and records indicate that some biological functions have been investigated more extensively than others, while linking activity annotations with chemical classification allows comparison of functional patterns across structurally distinct compound groups. Each activity record retains its source information and supporting literature where available, enabling provenance tracing. Importantly, the number of records associated with a compound or activity category reflects the amount of collected evidence rather than the magnitude or strength of the biological effect, supporting more appropriate compound-centered and activity-centered exploration.

**Fig. 3:**
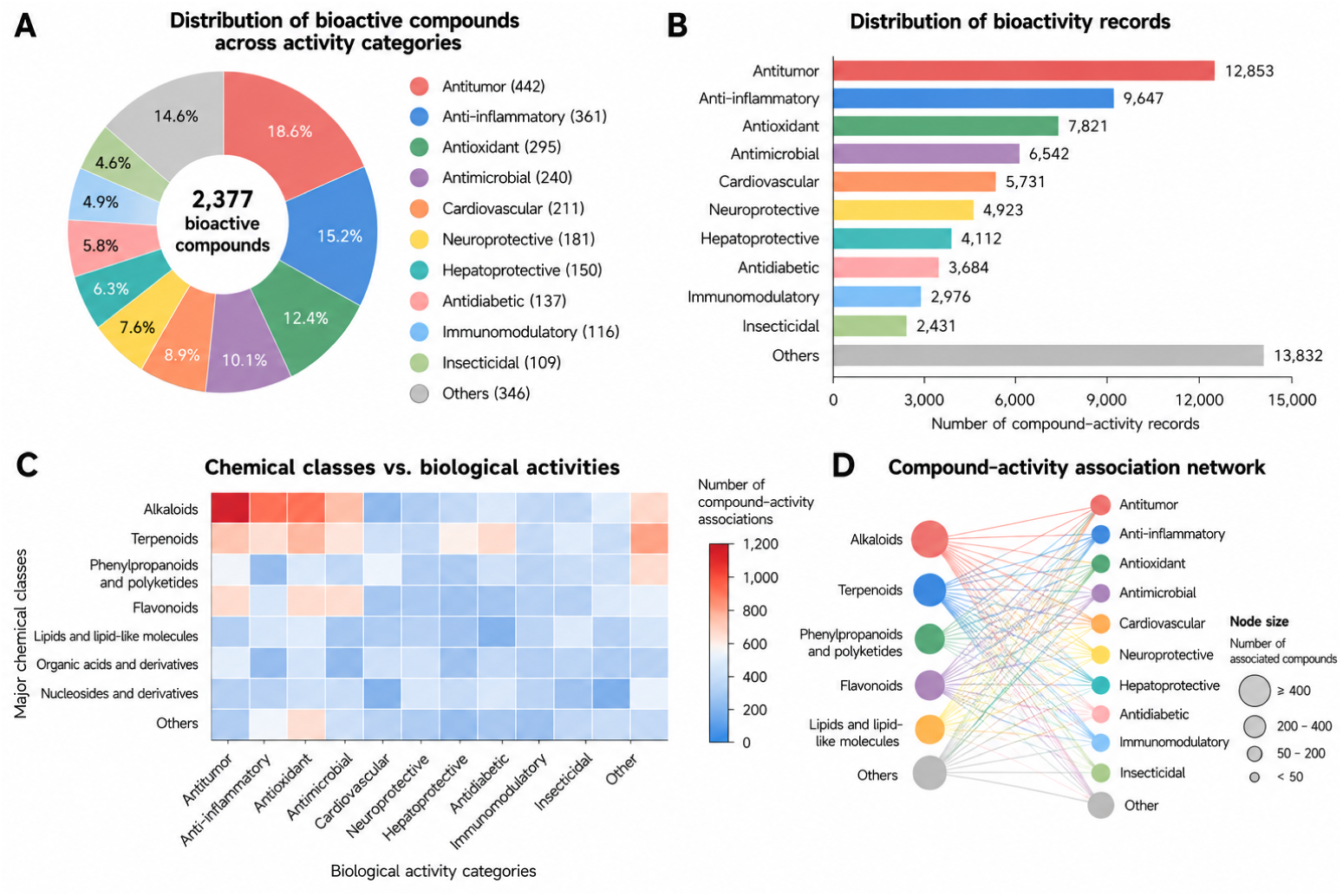
Landscape of biological activity annotations in TBACdb. (A) Distribution of the 2,377 bioactive compounds across major biological activity categories. (B) Distribution of the 64,522 compound–bioactivity records across major activity categories. (C) Heatmap showing associations between major chemical classes and biological activity categories. (D) Network representation of representative chemical class–activity associations, with node size indicating the number of associated compounds.

### 3.4 Species origins and literature distribution

To provide biological context for tobacco-associated compounds, TBACdb integrates species-origin information from natural-product databases, plant-oriented resources, and scientific literature. After compound identity matching, normalization, and record consolidation, 12,465 compound–species associations were retained, describing the reported occurrence of tobacco-associated chemical entities across diverse plant species. Because individual compounds may occur in multiple species and each species may contain multiple compounds, these relationships form a many-to-many network that helps users interpret compound distribution and biological activity annotations derived from non-tobacco plant resources. In parallel, TBACdb incorporates more than 3.7 million literature records linked to standardized compounds through compound names, synonyms, chemical identifiers, and related search terms. Species-origin, biological activity, and literature information can therefore be explored jointly within the same framework, allowing users to trace reported plant sources, functional annotations, and supporting publications. The overall distributions of species-origin and literature records are summarized in Fig. 4. Because the literature collection represents bibliographic associations rather than uniformly manually validated evidence, tobacco-focused publications are distinguished from broader literature associated with the same chemical entities.

**Fig. 4:**
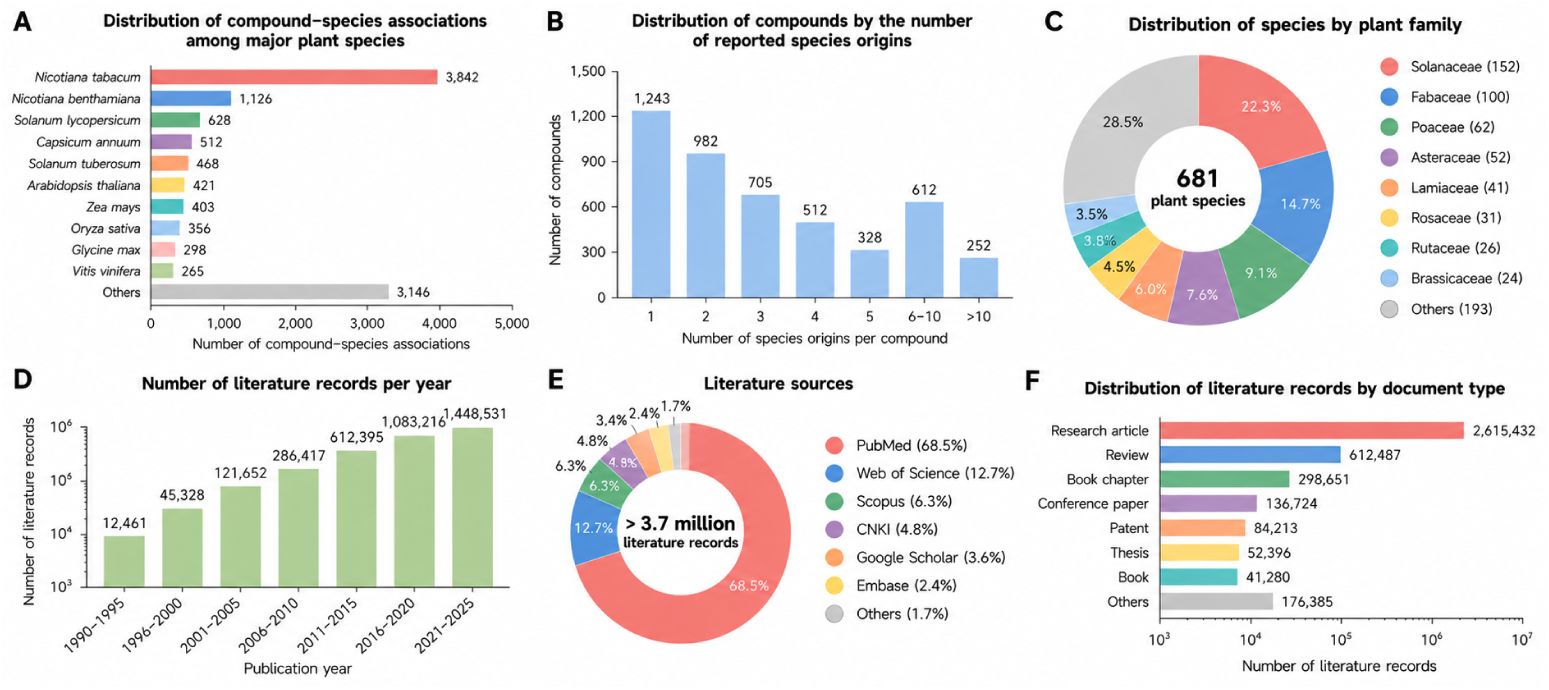
Overview of species-origin and literature information integrated in TBACdb. (A) Distribution of compound–species associations among major plant species. (B) Distribution of tobacco-associated compounds according to the number of reported species origins. (C) Distribution of recorded plant species across major taxonomic families. (D) Distribution of literature records across different publication periods. (E) Distribution of literature records according to major literature sources. (F) Distribution of literature records according to document type.

### 3.5 Coverage of physical reference standards

TBACdb currently links 783 compounds to corresponding physical reference standards, representing approximately 16.9% of the 4,634 standardized tobacco compounds included in the database. Each reference standard is connected to its digital compound record through a unified identifier, allowing users to simultaneously examine chemical structures and physicochemical properties, reported biological activities, species origins, associated literature, and the availability of experimentally accessible materials. Inventory information, including storage location, material status, and related management records, is maintained through the physical repository system to support traceability and resource management. This digital–physical linkage extends the role of TBACdb beyond information retrieval alone by providing a direct connection between candidate identification and downstream analytical work. Compounds with available standards can be more readily subjected to chemical confirmation, quantitative analysis, analytical method development, or subsequent functional validation. The overall coverage and organization of the physical reference repository are summarized in Fig. 5. Although the current standards represent only a subset of the total tobacco-associated chemical space, the repository provides a practical foundation for linking database-based discovery with experimental workflows and can be progressively expanded as additional compounds are prioritized and new standards become available.

**Fig. 5:**
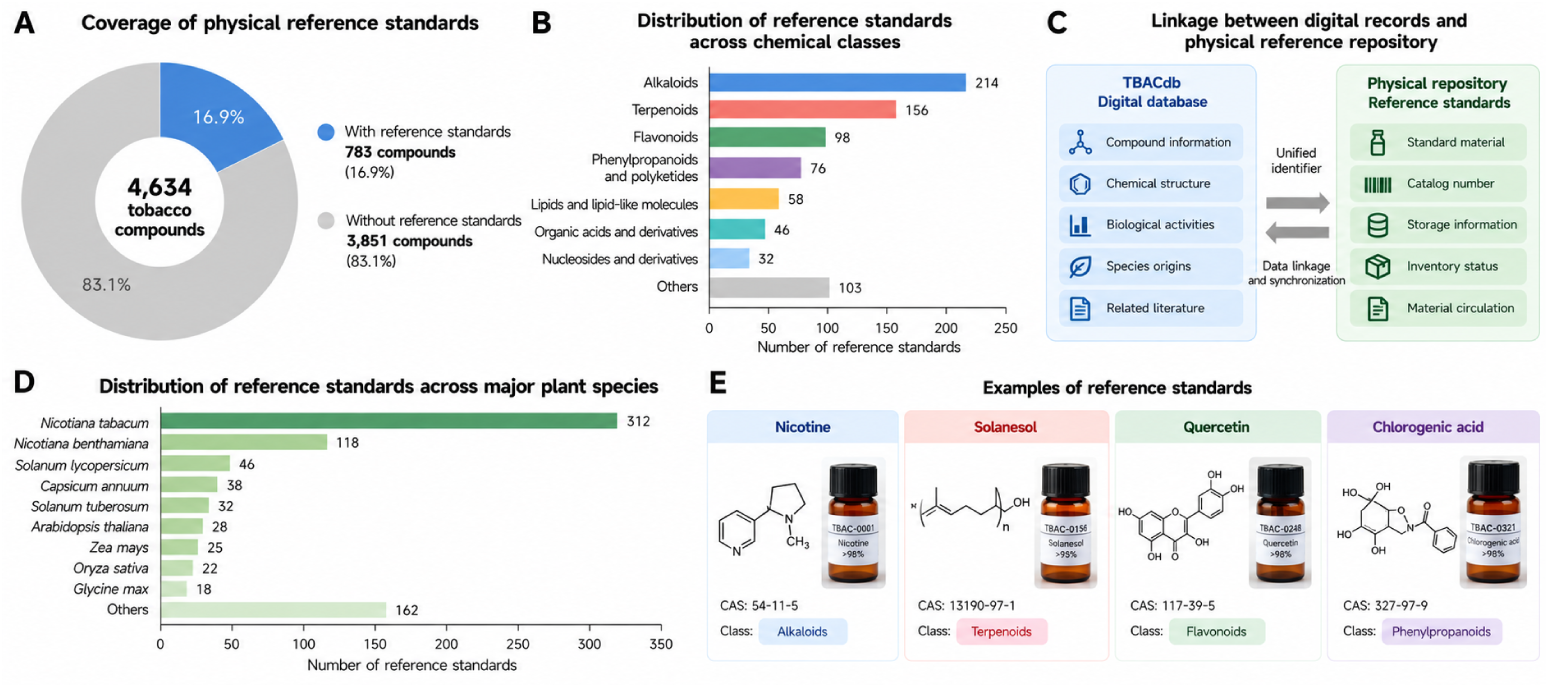
Coverage and organization of physical reference standards in TBACdb. (A) Coverage of physical reference standards among the 4,634 standardized tobacco compounds, including 783 compounds with available reference standards. (B) Distribution of physical reference standards across major chemical classes. (C) Schematic representation of the linkage and synchronization between digital compound records in TBACdb and the physical reference repository. (D) Distribution of compounds with available reference standards across representative plant species according to their recorded species origins. (E) Representative examples of physical reference standards and their corresponding digital compound records.

## 4 Database interface and functionality

### 4.1 Search and browse

TBACdb provides a web-based interface for searching and browsing tobacco-associated compounds through a unified entry point. Users can retrieve compounds using names, synonyms, CAS registry numbers, and related keywords, while browsing functions allow exploration without requiring an exact query. Search results provide basic compound information and direct links to detailed records, enabling users to move efficiently from an initial query to more comprehensive chemical and biological information. The interface is organized around standardized compound entities, so chemical properties, biological activities, species origins, literature associations, and physical reference-standard availability can be accessed within the same workflow. Users can therefore begin with either a known compound or a broader keyword and progressively explore related information without repeatedly searching independent databases. Where corresponding identifiers are available, links to external resources such as Pub- Chem, PubMed, and DrugBank are provided to facilitate access to complementary chemical, pharmacological, and bibliographic information.

### 4.2 Compound information and visualization

Each compound detail page presents multidimensional information associated with a standardized tobacco compound. Basic information includes compound name, CAS registry number, molecular formula, molecular weight, SMILES, InChI, InChIKey, physicochemical properties, chemical classification, and two-dimensional molecular structure where available. Biological activity annotations are displayed together with their corresponding source information and supporting literature, allowing users to examine multiple reported functions and evaluate their evidence context. Speciesorigin records provide additional information on the plant sources in which the same chemical entity has been reported and support exploration of compound–species relationships. Literature records are linked to individual compounds to facilitate evidence tracing, while external links allow users to extend their search to relevant chemical and biomedical databases. For compounds represented in the physical reference repository, the detail page also indicates the availability of corresponding standards and provides access to associated inventory information. By integrating chemical, biological, bibliographic, and physical-resource information in a single interface, TBACdb enables rapid interpretation of individual compounds and supports subsequent candidate selection and experimental planning.

### 4.3 Data access, sharing, and updates

TBACdb was designed as a continuously maintainable resource rather than a static database. Newly curated compounds, biological activity annotations, species-origin records, literature associations, and physical reference materials can be incorporated through the established data-processing and quality-control workflow. Version and provenance information are retained during database maintenance to support record tracing, correction, and future updating. The digital database and physical reference repository are managed in coordination, allowing changes in reference-standard inventory, storage status, and circulation information to be reflected in the corresponding compound records. The platform also supports statistical analysis, authorized data sharing, and resource distribution under the relevant management requirements. Together with external database linkage and continuous data updating, these functions facilitate the long-term maintenance, reuse, and expansion of TBACdb and provide a practical framework for both information retrieval and resource-oriented tobacco bioactive compound research.

## 5 Case studies

To illustrate the practical utility of TBACdb, we considered three representative usage scenarios covering compound-centered exploration, activity-oriented candidate screening, and linkage between digital information and physical reference materials. These scenarios represent common workflows in tobacco natural-product research, in which users may begin with a known compound, a biological function of interest, or a set of candidates requiring subsequent experimental verification. Together, they demonstrate how the chemical, biological, bibliographic, and physical-resource layers of TBACdb can be combined within a unified workflow (Fig. 6).

**Fig. 6:**
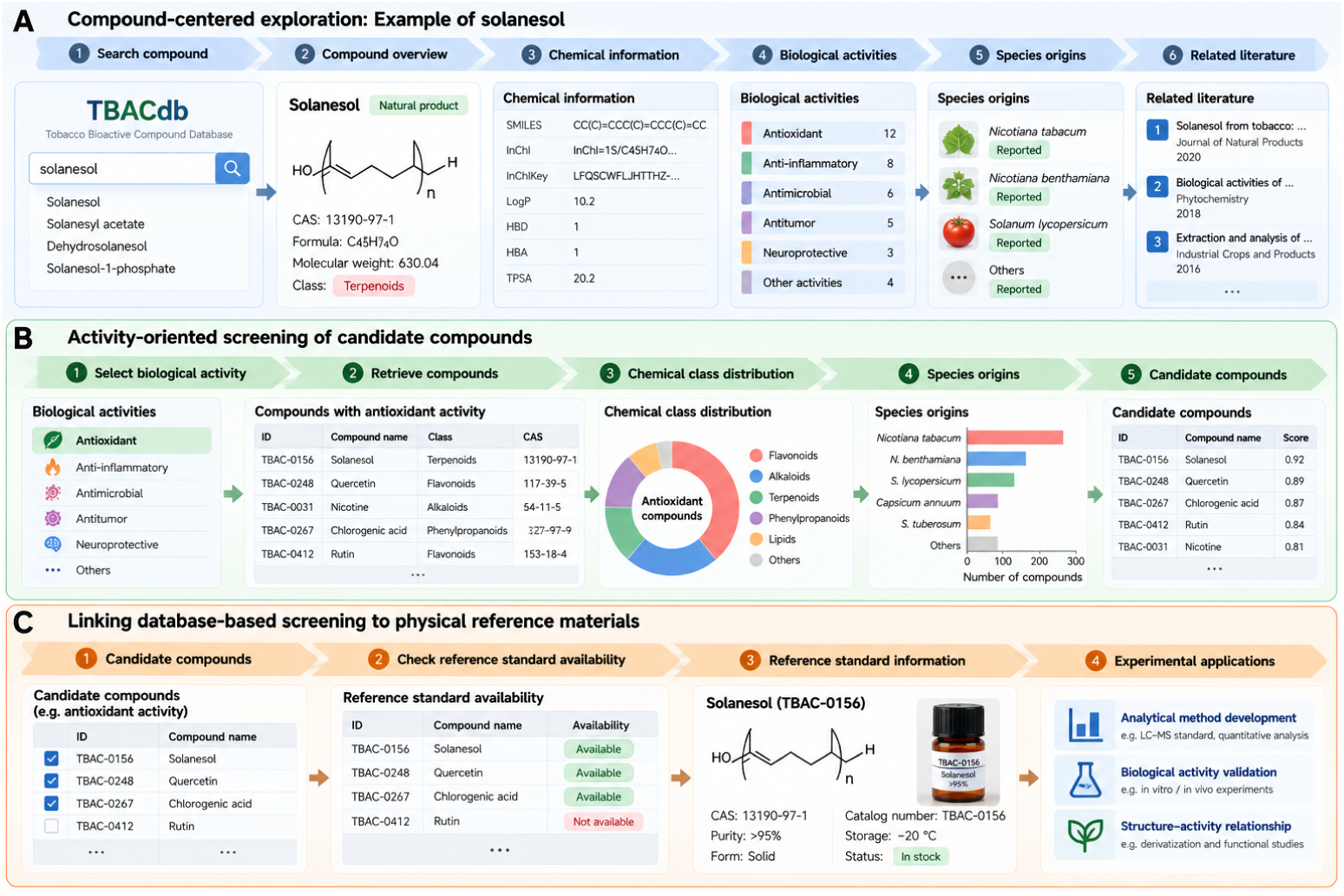
Representative applications of TBACdb. (A) Compound-centered exploration using solanesol as an example, showing the workflow from compound search to chemical information, biological activities, species origins, and related literature. (B) Activity-oriented screening of candidate compounds, illustrating retrieval of compounds associated with a selected biological activity and subsequent comparison based on chemical class, species origins, and candidate prioritization. (C) Linking databasebased candidate screening to the physical reference-material repository, enabling users to determine reference-standard availability and support downstream analytical and experimental applications.

### 5.1 Compound-centered exploration of solanesol

Solanesol is a representative tobacco-associated natural product that has been widely investigated for its biological activities and potential applications [7]. It was therefore selected to demonstrate compound-centered exploration in TBACdb. By entering “solanesol” or its CAS registry number into the search interface, users can directly access the corresponding standardized compound record. The detail page integrates basic chemical information, including molecular formula, molecular weight, twodimensional structure, SMILES, InChI, InChIKey, physicochemical properties, and chemical classification. From the same record, users can further inspect reported biological activities, identify plant species in which solanesol has been reported, and retrieve associated literature. The source information retained for individual annotations also allows users to examine the evidence context of reported activities rather than treating all functional records as equivalent. External links to PubChem and PubMed provide access to complementary chemical and bibliographic information when additional detail is required. If a corresponding physical reference standard is available, its availability can also be identified through the same compound record. This example illustrates how TBACdb can consolidate information that would otherwise require searches across several independent chemical, natural-product, and literature databases and provides a convenient starting point for subsequent analytical or functional studies.

### 5.2 Activity-oriented screening of candidate compounds

TBACdb can also be explored from a biological activity perspective when a research question begins with a desired function rather than a predefined compound. For example, users interested in antioxidant, anti-inflammatory, antimicrobial, antitumor, or other reported activities can retrieve tobacco-associated compounds linked to the corresponding activity annotations. Candidate compounds can then be compared according to their chemical classes, species origins, associated literature, and other available annotations. Because individual compounds may be linked to multiple biological activities, this workflow also allows users to identify compounds with overlapping functional annotations and to explore whether particular chemical classes are repeatedly associated with a selected activity category. Importantly, TBACdb retains the provenance of activity records. Users can therefore distinguish evidence derived from tobacco-focused studies from broader biological information reported for the same chemical entity in other plant-derived resources. This evidence-aware screening strategy reduces the risk of interpreting cross-plant annotations as direct tobacco-specific validation and provides a more appropriate basis for narrowing a large compound collection to a smaller set of candidates for subsequent experimental investigation.

### 5.3 Linking database-based screening to physical reference materials

A further application of TBACdb is the translation of database-based candidate prioritization into experimentally accessible resources. After compounds of interest have been identified according to chemical properties, biological activities, species origins, or literature evidence, users can determine whether corresponding physical reference standards are available in the integrated repository. For compounds with available standards, the standardized digital record is linked to the physical-resource record through a unified identifier, allowing users to access chemical information, biological evidence, supporting literature, and material availability within the same workflow. This digital–physical connection is particularly useful when several candidate compounds satisfy similar biological criteria but differ in experimental accessibility. Availability of a reference standard can facilitate subsequent compound identification, quantitative analysis, analytical method development, and functional validation, whereas compounds without available standards may be considered priorities for future expansion of the repository. The integration of database-based screening with physical-material information therefore provides a practical route from computational or literature-based candidate identification to downstream laboratory investigation.

Together, these case studies illustrate three complementary modes of using TBACdb: starting from a known tobacco compound, starting from a biological activity of interest, and translating candidate identification into experimentally accessible materials. These workflows demonstrate how TBACdb can bridge multidimensional information integration, evidence-aware candidate prioritization, and subsequent analytical and experimental investigation.

## 6 Discussion

TBACdb was developed to address the fragmentation of chemical, biological, and literature information related to tobacco-associated compounds. Recent natural-product databases, including NPBS Atlas, Natural Products Atlas 3.0, and NPASS 3.0, have substantially expanded the integration of chemical structures, species origins, and biological activities [24–26]. However, these resources focus on broader natural-product collections or specific biological domains rather than tobacco-associated compounds. TBACdb complements these databases by integrating standardized tobacco compounds, biological activities, species origins, literature evidence, and physical reference materials within a unified compound-centered framework.

An important feature of TBACdb is the integration of heterogeneous biological information while preserving evidence provenance. Standardized chemical identifiers are essential for reconciling compound records across different resources, as demonstrated by recent metabolite data harmonization approaches [27]. Tobaccofocused literature curation identified 166 compounds with directly reported biological activities, while cross-resource integration expanded functional coverage to 2,377 compounds. Importantly, biological activities reported for the same chemical entities in other plant resources should not be interpreted as direct experimental validation in tobacco-derived materials. Retaining source information therefore supports more reliable interpretation and candidate prioritization.

Another distinctive feature is the linkage between digital compound records and 783 physical reference standards, representing approximately 16.9% of the standardized compound collection. Reference materials are important for improving analytical accuracy, reproducibility, and quantitative consistency in metabolomics research [28, 29]. This digital–physical integration provides a practical connection between database-based compound screening and subsequent analytical verification, method development, and experimental investigation.

Several limitations remain. Biological activity records originate from heterogeneous sources with varying levels of experimental support, while the extensive literature collection represents bibliographic associations rather than uniformly validated evidence. In addition, the physical repository currently covers only a subset of the database compounds. Future development will focus on evidence grading, quantitative bioactivity annotation, molecular target and pathway integration, and expansion of referencestandard coverage. Such improvements may further support the identification and experimental investigation of biologically relevant natural products [30, 31].

## 7 Conclusions

TBACdb provides an integrated resource for tobacco-associated bioactive compounds, containing 4,634 standardized compounds, 2,377 compounds with reported biological activities, and 783 linked physical reference standards. By integrating chemical information, biological activities, species origins, literature evidence, and physical materials within a unified compound-centered framework, TBACdb supports compound retrieval, candidate screening, and subsequent experimental investigation. Continued data curation and expansion of the physical repository will further enhance its value for tobacco bioactive compound research.

## Supporting information

sn-article.tex

