## Supplementary figures and images for "TBACdb: a comprehensive database of tobacco bioactive compounds"

### fig1.png

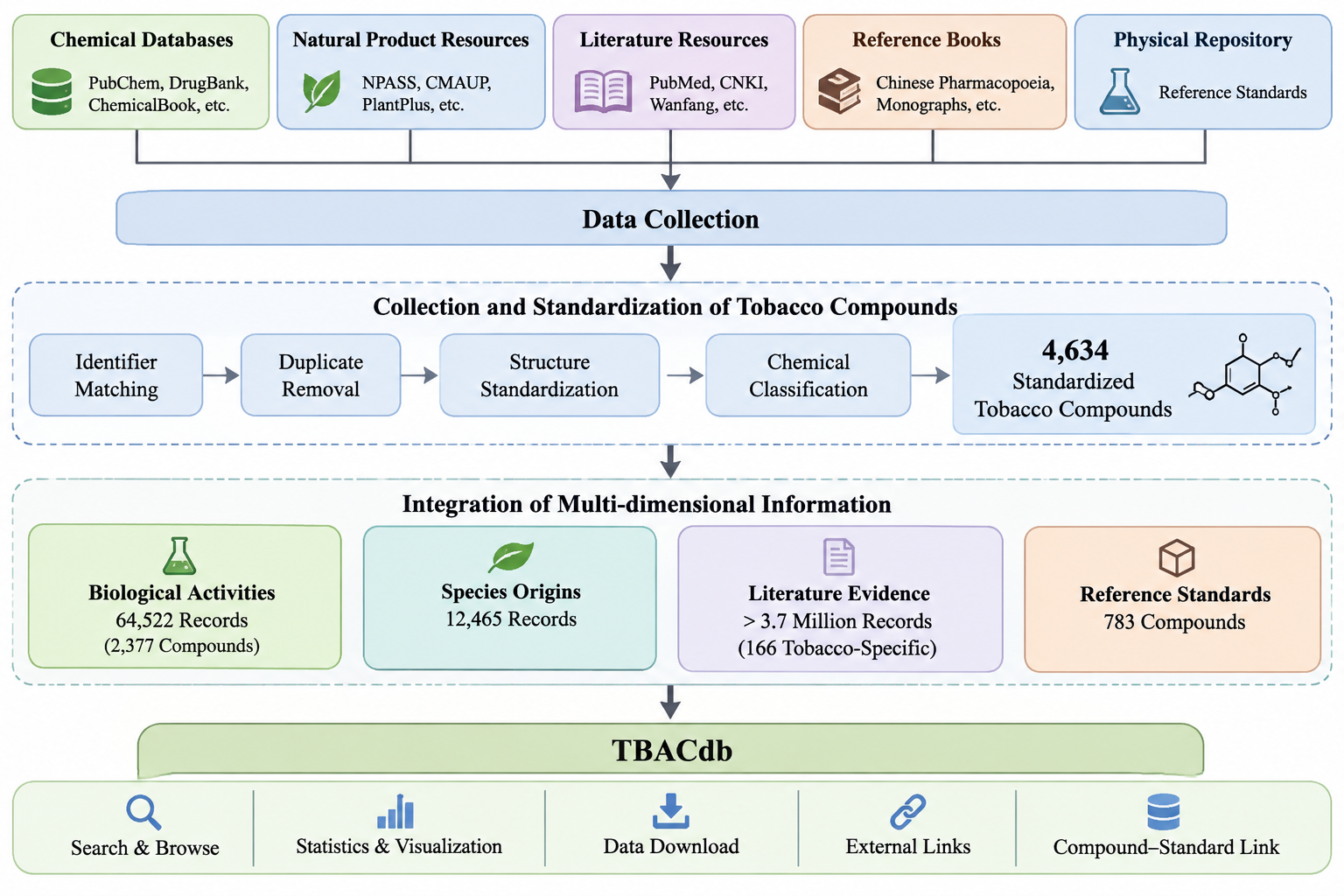

### fig2.png

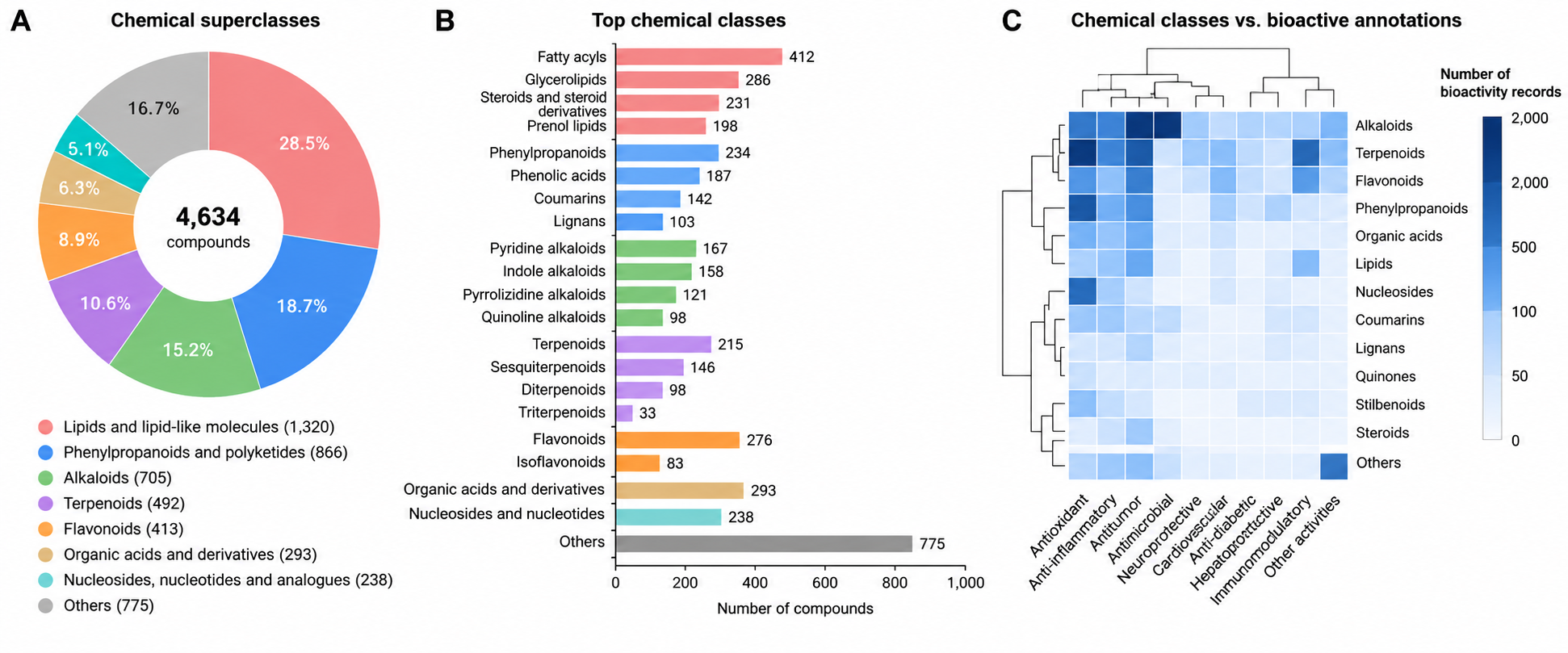

### fig3.png

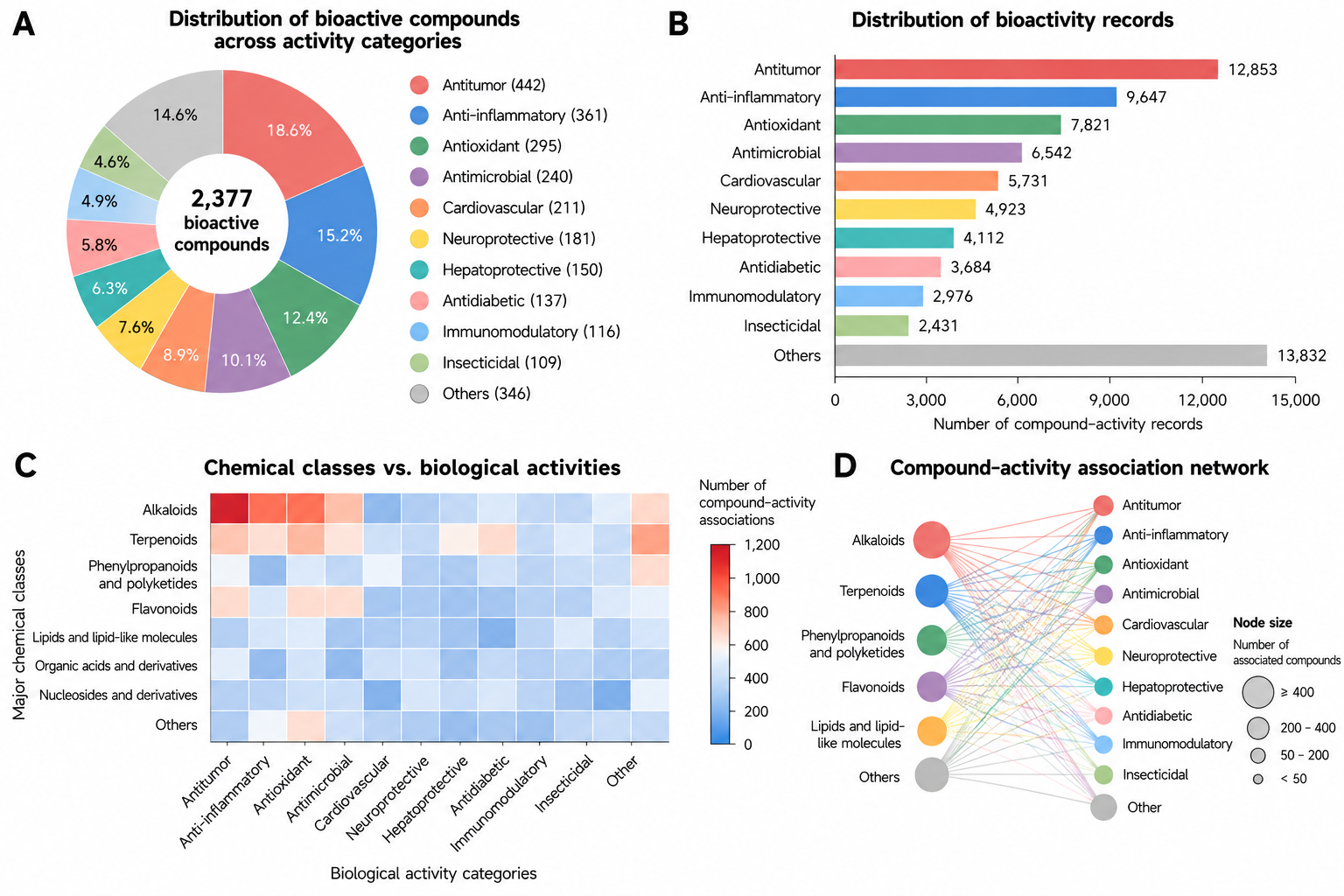

### fig4.png

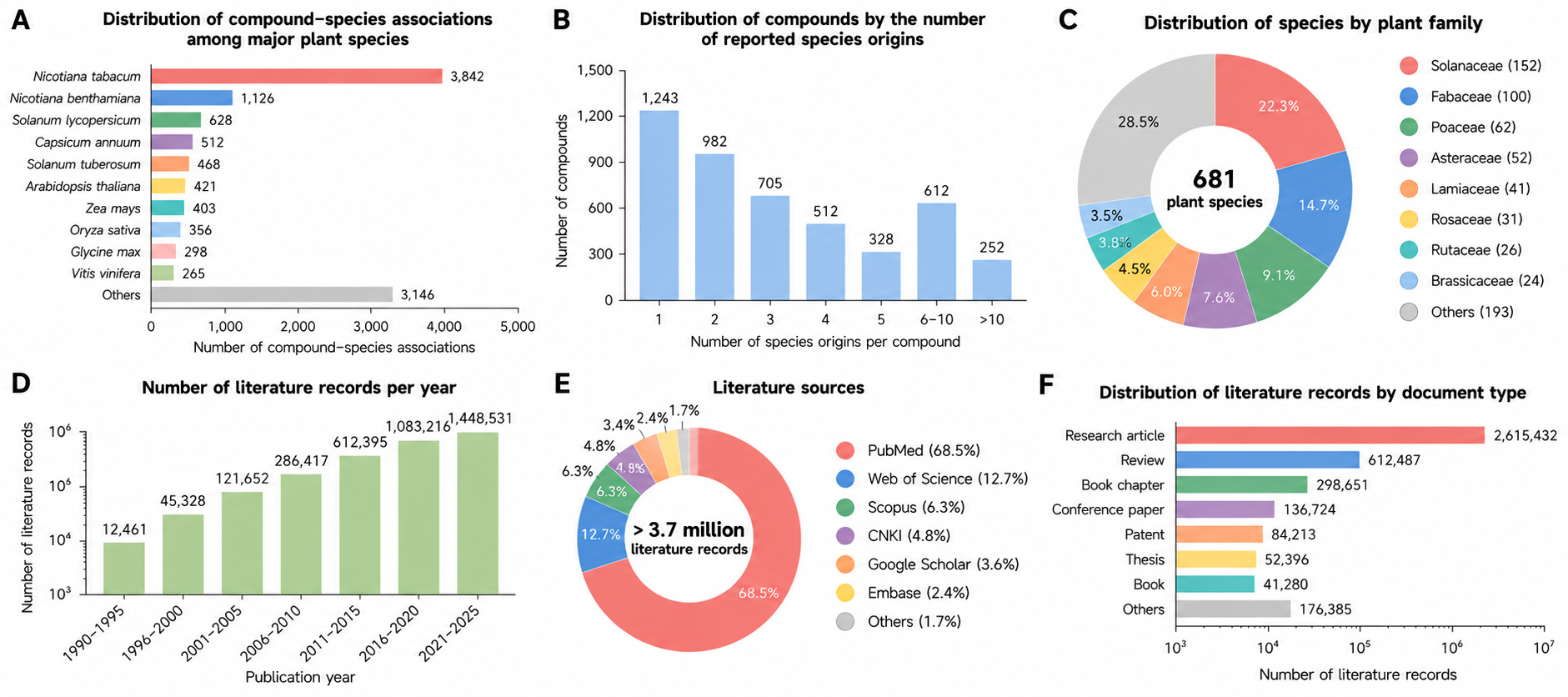

### fig5.png

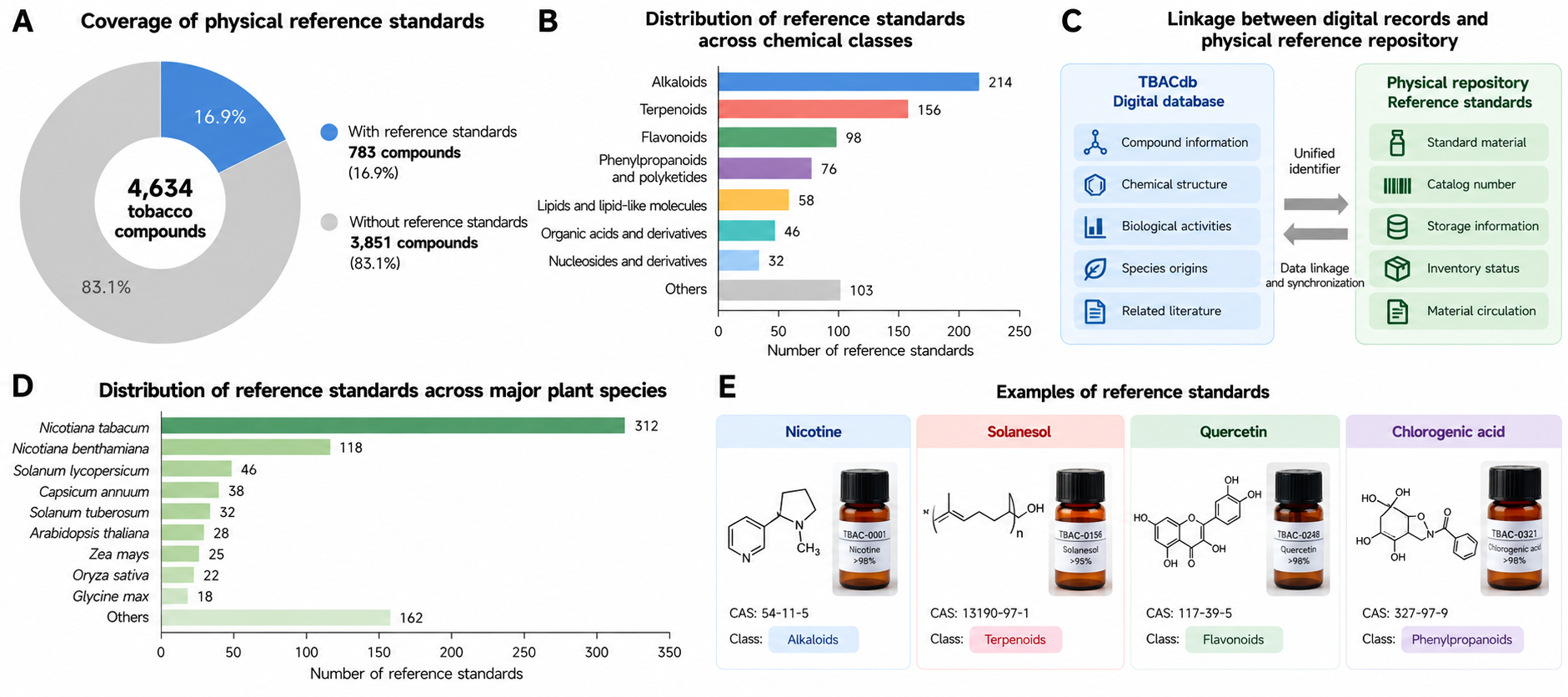

### fig6.png

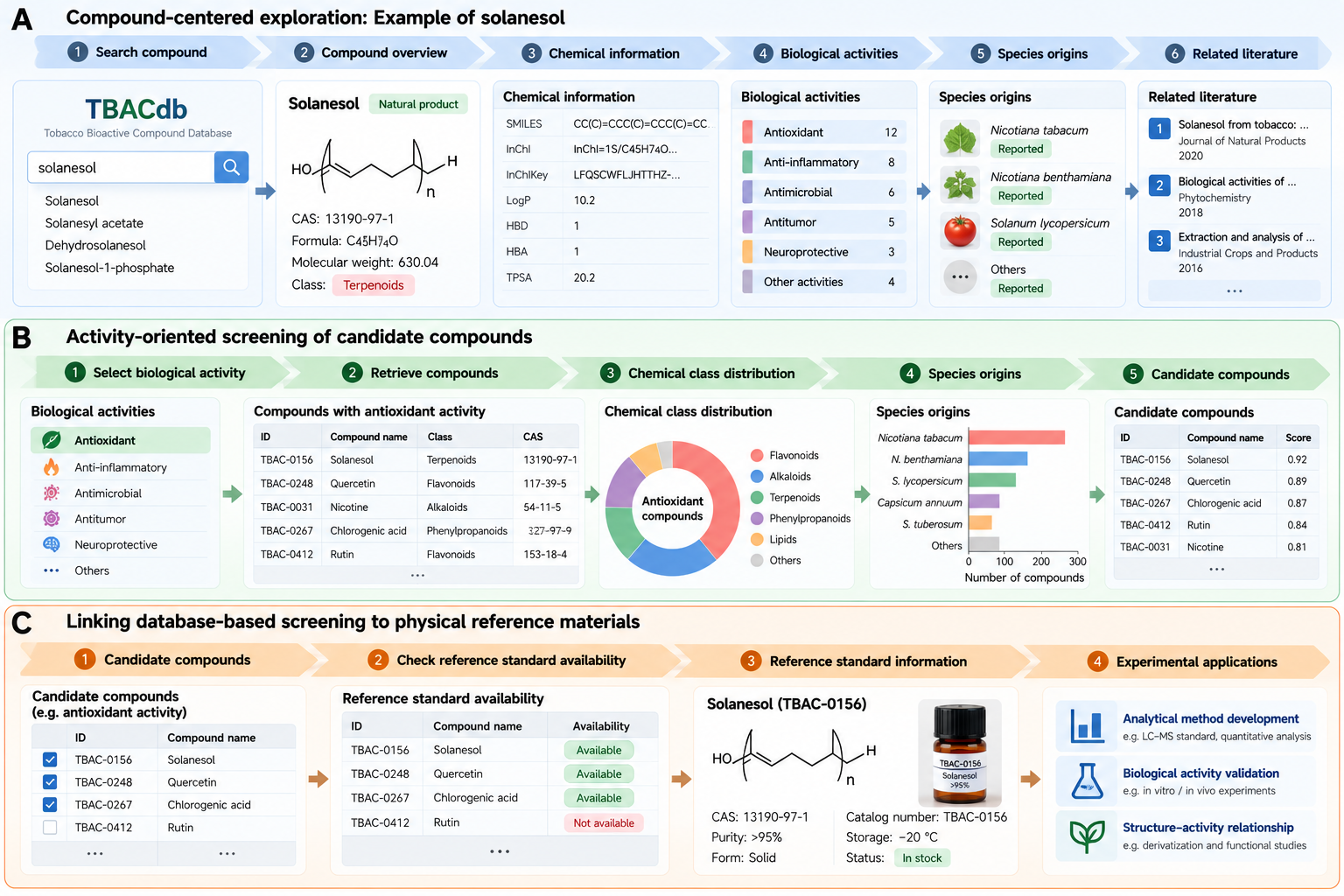
